# Cyclin D1·Cdk4 regulates neuronal activity through phosphorylation of GABA_A_ receptors

**DOI:** 10.1101/2022.06.30.498219

**Authors:** Neus Pedraza, Ma Ventura Monserrat, Francisco Ferrezuelo, Jordi Torres-Rosell, Neus Colomina, David Soto, Federico Miguez-Cabello, José A. Esteban, Joaquim Egea, Eloi Garí

## Abstract

Cyclin D1 (Ccnd1)·Cdk4 complexes drive cell cycle progression through phosphorylation of pRb. Interestingly, Ccnd1 moves to the cytoplasm at the onset of differentiation in neuronal precursors. However, the cytoplasmic functions and targets of Ccnd1 in post-mitotic neurons are unknown. Here we identify the α4 subunit of gamma-aminobutyric acid (GABA) type A receptors (GABA_A_Rs) as an interactor and target of Ccnd1·Cdk4. Ccnd1 binds to an intracellular loop in α4 and, together with Cdk4, phosphorylates the α4 subunit at threonine 423 and serine 431. These modifications increase the activity of α4-containing GABA_A_Rs, measured in whole-cell patch-clamp recordings, and upregulate its surface levels. In agreement with this role of Ccnd1·Cdk4 in neuronal signaling, inhibition of Cdk4 decreases synaptic and extrasynaptic currents in the hippocampus of newborn rats. Moreover, *CCND1* knockout mice display an altered pattern of dendritic spines, according to α4 functions in synaptic pruning. Overall, our findings molecularly link Ccnd1·Cdk4 to GABA_A_Rs activity in the central nervous system and highlight a novel role for this G1 cyclin in neuronal signaling.

## INTRODUCTION

Cyclin D1 (Ccnd1) and its catalytic counterparts, cyclin-dependent kinase 4 (Cdk4) and Cdk6, are main regulators of cell proliferation by promoting the G1-S phase transition of the cell cycle (reviewed in[1]). Under favorable conditions, mitogenic signals induce the expression of Ccnd1, the formation of Ccnd1·Cdk4/6 complexes and their localization to the nucleus. There, they phosphorylate the Retinoblastoma repressor protein (pRB), which eventually leads to its inhibition and in turn releases the activity of the E2F transcription factor. This induces the expression of genes required to go through the next steps of the cell cycle. This pRB-dependent control of cell proliferation is considered the “canonical” function of Ccnd1 and it has been reported as very relevant in tumor proliferation. Accordingly, Ccnd1 is classified as an oncogene frequently amplified in many types of neoplasia, and different cancer treatments with Cdk4/6 inhibitors such as Palbociclib have been approved to block this function. However, Ccnd1 also operates through several other non-canonical pathways including some pRB- and Cdk-independent ones to regulate different cellular processes (reviewed in[2]). Thus, Ccnd1 promotes cell detachment, migration and invasion of normal and tumor cells independently of pRB status[3],[4],[5] by interacting with cytoplasmic and membranous targets such as filamin A, pacsin, RalGTPases and paxillin[6],[7],[8],[9]. For example, Ccnd1·Cdk4 promotes cell invasion and metastasis through the phosphorylation of paxillin and the activation of the Rac1 pathway in membrane ruffles[9]. Moreover, the expression of a membrane-associated form of Ccnd1 harboring the pharnesylation signal of K-Ras (Ccnd1-CAAX) maximizes the ability of tumor cells to invade and metastasize[10]. Besides cell invasion, membrane-targeted Ccnd1 also influences cell-signaling responses[11].

The first hint for a role of Ccnd1 in the nervous system came from the analysis of *CCND1* knockout (KO) mice[12], which present certain behaviors (altered leg-clasping reflex) indicative of neurological abnormalities. Since then, several studies have substantiated the involvement of Ccnd1 in neuronal functioning. In neural stem cells, Ccnd1·Cdk4 has been involved in regulating the length of G1, which is thought to influence the decision of cells to proliferate or differentiate[13],[14],[15]. Also, under pathological conditions, nuclear Ccnd1 reactivation/ upregulation in post-mitotic neurons has been observed to promote apoptosis (reviewed in[2]). Ccnd1 is located in the cytoplasm of differentiating post-mitotic cortical neurons and in neuroblastoma cell lines, although this has been proposed as a mechanism to prevent apoptosis[16] or to promote cell cycle withdrawal[17], respectively. Nevertheless, Ccnd1 could be exerting an active role in the cytoplasm. Interestingly, Ccnd1 is necessary for NGF-induced neuritogenesis in PC12 pheochromocytoma cell line[18]. Also cytoplasmic expression of Ccnd1·Cdk4 has been observed in the hippocampus during development[19] where it has been involved in neuronal plasticity[20]. But despite these works, the role and target/s of Ccnd1 in the nervous system and in particular its role in the cytoplasm of the neurons remains poorly understood.

γ-aminobutyric acid (GABA) is the major inhibitor neurotransmitter in the central nervous system. Its actions are mediated through different types of GABA receptors: GABA A receptors (GABA_A_Rs), which are ligand-gated chloride ion channels, and GABA B receptors (metabotropic G-protein-coupled). Deficits in GABA_A_R function are increasingly being involved in different pathologies, such as anxiety, cognitive deficits, depression, epilepsy, schizophrenia and substance abuse[21]. These receptors are also clinically relevant drug targets for anesthetic, anticonvulsant, anxiolytic, or sedative-hypnotic agents. GABA_A_Rs mediate both phasic and tonic inhibition: synaptic GABA_A_Rs are activated transiently by the GABA released from presynaptic vesicles, whereas extrasynaptic GABA_A_Rs are activated by low GABA concentrations in the extracellular space and thus mediate tonic inhibition. The tonic inhibitory currents control neuronal excitability and the strength of synaptic transmission[22].

GABA_A_Rs are heteropentamers that belong to the ligand-gated ion channel superfamily. A total of 19 subunits (as well as splice variants) have been cloned and grouped into eight subunit classes: α (1-6), β (1-3), γ (1-3), δ, ε, π, θ and ρ (1-3). GABA_A_Rs are usually formed by two α, two β, and one γ or one δ subunits. Each subunit contains a large amino-terminal extracellular domain and four transmembrane (TM) domains. Between TM3 and TM4 there is a large intracellular loop (ICL) where regulation by phosphorylation and protein interactions primarily occurs. The subunit composition dictates its subcellular location and function.

GABA_A_Rs containing the α4 subunit are expressed in the hippocampus, cortex and thalamus, and mediate extrasynaptic inhibition in thalamus and dentate gyrus[23]. At the onset of puberty, α4-β-δ GABA_A_R expression increases perisynaptically at excitatory synapses in the hippocampus, contributing to anxiety, synaptic pruning and learning deficits in mice[24],[25],[26],[27],[28]. In fact, in the α4-KO mice synaptic pruning is prevented, and synaptic plasticity and learning abilities are restored[23],[29],[30]. In the adult hippocampal neurogenesis, α4-containing GABA_A_Rs are involved in proliferation, migration and dendritic development[31]. Mutations in the GABAergic receptor subunit 4 gene *GABRA4*[32],[33],[34] and reduced protein and mRNA levels of α4[35],[36] are reported in patients of autism spectrum disorder. Furthermore, the α4-KO mice show a phenotype compatible with high-functioning autism[37].

Here we report that Ccnd1 interacts with the α4 subunit of GABA_A_Rs. Ccnd1·Cdk4 complex phosphorylates α4 in the intracellular loop between TM3 and TM4 and enhances the efficacy of α4-containing GABA_A_Rs, affecting the pattern of dendritic spines. We propose a novel role for cytoplasmic Ccnd1·Cdk4/6 in regulating α4-containing GABA_A_Rs in the central nervous system.

## RESULTS

### The α4 subunit of GABA_A_ receptors interacts with cyclin D1

To investigate the role of Ccnd1 in the nervous system, we performed a yeast two-hybrid screening using Ccnd1 as a bait and an adult mouse brain library as a prey[38]. We found that Ccnd1 interacts with the C-terminal half of the α4 GABA_A_R subunit (Gabra4). We obtained three independent clones spanning amino acids 354-552, 385-552 and 386-552 (Fig 1A). This region of α4 contains the intracellular loop (ICL) between the TM3 and TM4 transmembrane domains, which is phosphorylated by protein kinase C (PKC,[39]). To confirm the interaction between the α4 subunit and Ccnd1 we carried out immunoprecipitation (IP) experiments with FLAG-tagged Ccnd1 and the HA-tagged C-terminal region of α4 (354-552aa) expressed in HEK-293T cells. We observed specific co-IP of the C-terminal domain of α4 in the Ccnd1 IP (Fig 1B). To test whether there is a direct interaction between Ccnd1 and the α4 subunit, we performed *in vitro* GST-pull-down assays. Fusions of GST-α4 or GST-C-terminal α4 (354-552aa) purified from *E.coli* were mixed with FLAG-Ccnd1 produced by *in vitro* transcription and translation. Ccnd1 was co-precipitated when GST-α4 or GST-C-terminal α4 were pulled down, but not with GST alone (Fig 1C). To analyze the endogenous interaction between Ccnd1 and the α4 GABA_A_R subunit, we immunoprecipitated Ccnd1 from the hippocampus of adult mice, since co-expression of Ccnd1 and α4 mRNAs in the CA1, CA3 and dentate gyrus regions of the hippocampus is observed in the Allen mouse brain atlas (www.brain-map.org). As shown in figure 1D, α4 subunit of GABA_A_R co-immunoprecipitated with endogenous Ccnd1, in a specific manner. Overall, our data indicate that there is a specific and direct interaction between Ccnd1 and the α4 subunit of GABA_A_ receptors.

**Figure 1.**
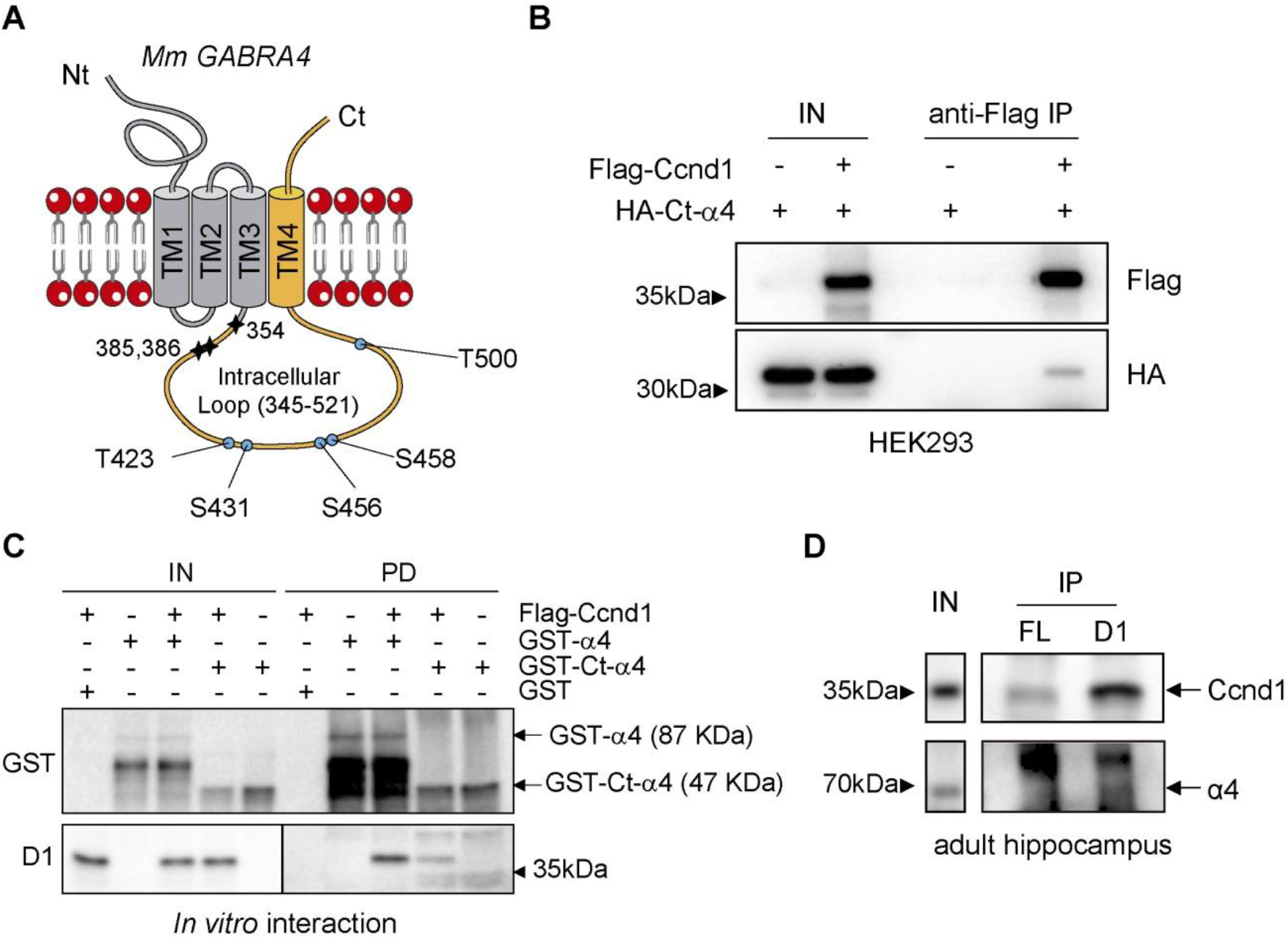
Ccnd1 interacts with the α4 subunit of GABA_A_ receptors. A) Scheme representing the mouse α4 subunit of GABA_A_Rs (552aa). N terminal (Nt) and C terminal (Ct), intracellular and extracellular domains and the four transmembrane domains (TM1-4) are shown. In a yeast two-hybrid screening, Ccnd1 was found to interact with three independent clones of the C-terminal region of the α4 subunit (in yellow), spanning amino acids 354-552, 385-552, 386-552 (the starting amino acids are shown as stars). The three clones contained the intracellular loop (ICL) between TM3 and TM4. Putative phosphorylation sites for Cdk (SP or TP) in the ICL are shown in blue. B) HEK-293T cells were transfected with the C terminus of α4 subunit (354-552aa, tagged with HA: HA-Ct-α4) and Ccnd1 (tagged with Flag: Flag-Ccnd1). Flag-Ccnd1 was precipitated using αFlag agarose beads (Sigma). Input (IN) and immunoprecipitation (IP) samples were analyzed by western blot using anti-HA and anti-Flag antibodies, to detect co-IP of the C terminus of α4 with Ccnd1. C) In vitro translated FLAG-Ccnd1 was incubated with recombinant GST or GST fusions of α4 full length (GST-α4) or the C terminus of α4 (GST-Ct-α4) purified from E. coli. Input (IN) and GST pull down (PD) samples were analyzed by western blot using anti-Flag antibody to detect Ccnd1 and anti-GST antibody to detect α4. Intact protein fusions are indicated. D) Immunoprecipitation of endogenous Ccnd1 from adult mouse hippocampal extracts was performed using a rabbit polyclonal antibody against Ccnd1. As a control, a rabbit polyclonal antibody against the Flag epitope (FL) was used. Input (IN) and IP samples were analyzed by western blot to detect Ccnd1 and α4 subunit.

### The α4 subunit of GABA_A_Rs is phosphorylated at T423 and S431 by Ccnd1·Cdk4 complex

As Ccnd1 is the regulatory subunit of Cdk4/6 kinases, we analyzed the sequence of α4 for putative phosphorylation sites by Cdks. We found five S/TP sites in the sequence of the intracellular loop (Fig 1A). In an *in vitro* kinase assay, Ccnd1·Cdk4 complex phosphorylated recombinant GST-α4 and this phosphorylation was prevented by inhibiting the complex with the specific Cdk4/6 inhibitor Palbociclib (Fig 2A). Besides, another subunit of the GABA_A_Rs, β3, which lacks S/TP sites, was not phosphorylated by the Ccnd1·Cdk4 complex (Fig 2B). In order to know which residues of α4 were phosphorylated by Ccnd1·Cdk4, we mutated the five intracellular putative Cdk phosphorylation sites to non-phosphorylatable residues (alanine) generating three different mutant alleles: T423A and S431A (α4^T423AS431A^); S456A and S458A (α4^S456AS458A^); and T500A (α4^T500A^). Neither α4^S456AS458A^ nor α4^T500A^ mutants showed a decrease in phosphorylation by Ccnd1·Cdk4 (Fig 2C and 2D). However, the non-phosphorylatable mutant of T423 and S431 (α4^T423AS431A^: AA) significantly reduced phosphorylation by Ccnd1·Cdk4 *in vitro*, either with the full-length or the C-terminus construct of α4 (Fig 2C-F). Moreover, we analyzed by mass spectrometry phosphopeptides from the C-terminus of α4 subunit phosphorylated by Ccnd1·Cdk4, and we found that both T423 and S431 were phosphorylated *in vitro* (S1 Table). Altogether, our results show that Ccnd1·Cdk4 complex is able to phosphorylate α4 subunit at T423 and S431.

**Figure 2.**
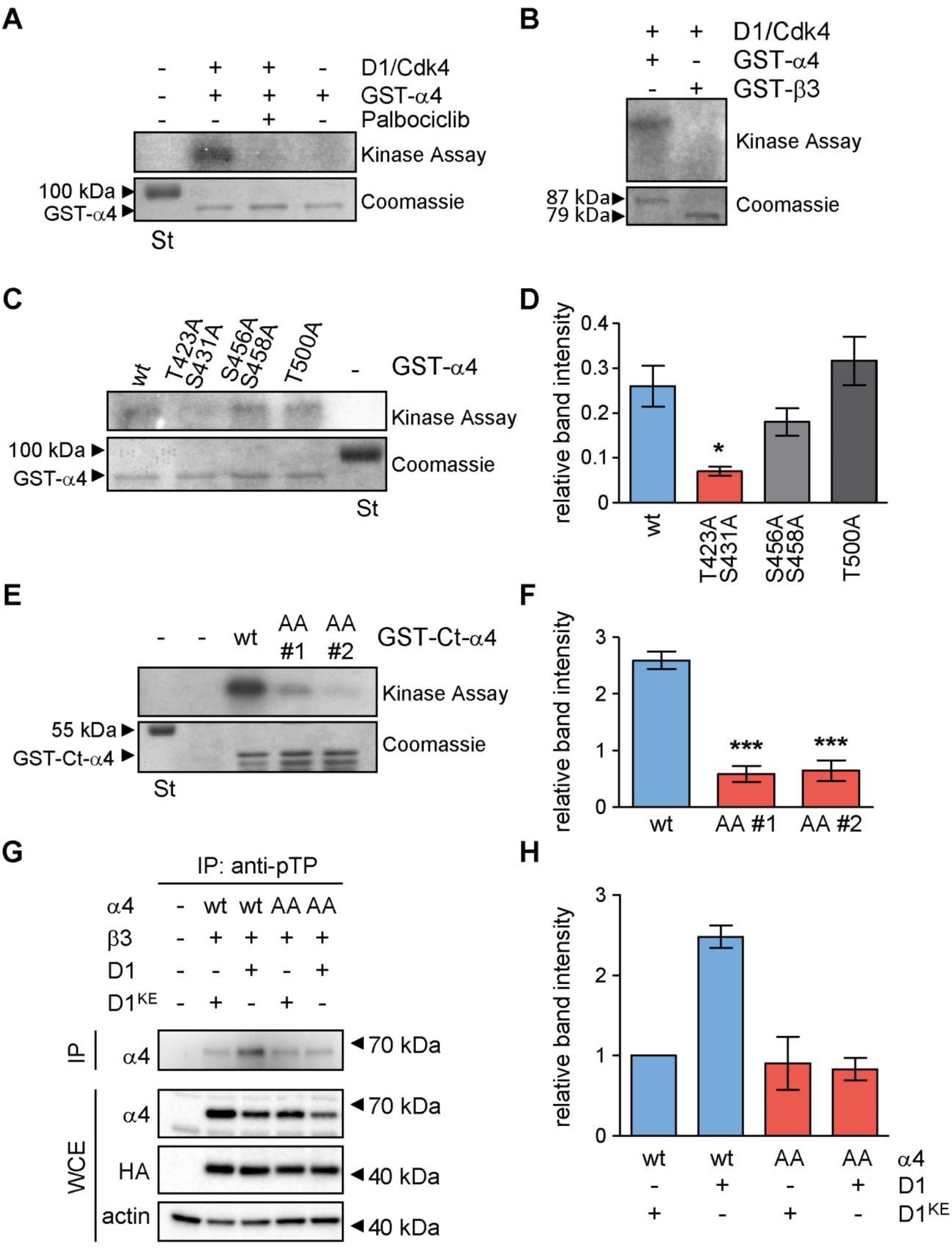
Ccnd1·Cdk4 phosphorylates α4 at T423 and S431 residues. A) In vitro radioactive kinase assay using purified recombinant GST-α4 fusion and Ccnd1·Cdk4 (Sigma). Cdk4/6 inhibitor Palbociclib was added at 1 μM concentration. Coomassie blue staining is shown as loading control (Input). B) Recombinant GST-α4 and GST-β3 fusions were tested for Ccnd1·Cdk4 kinase activity as in A. C) Kinase assay was performed with Ccnd1·Cdk4 against GST-α4 wild type and the non phosphorylatable mutants: T423A, S431A; S456A, S458A; and T500A. D) Quantification of the phosphorylation in C showing the mean ± S.E.M. The amount is relative to a pRB control (n=3, One-way ANOVA, post-Tukey analysis *p<0.05). E) Kinase assay was performed with Ccnd1·Cdk4 against the C-terminus of α4 wild type (GST-Ct-α4 wt) and two clones (#1 and #2) of the non phosphorylatable mutant T423A, S431A (AA). F) Quantification of the phosphorylation in E showing the mean ± S.E.M. The amount is relative to a pRB control (n=3, ANOVA ***p<0.001). G) Immunoprecipitation of Cdk substrates in HEK-293T cells transfected with α4 wild type (wt) or the non phosphorylatable mutant T423A, S431A (AA), together with β3 and HA-Ccnd1-CAAX (D1) or HA-Ccnd1^K112E^-CAAX (D1^KE^). PhosphoTP antibody was used for the immunoprecipitation (IP), and α4 was detected by immunoblot. Protein levels of α4, Ccnd1 (anti-HA antibody) and actin were also analyzed in the whole cell extract (WCE). H) Quantification of the phosphorylation in G showing the mean ± SD (n=2).

To investigate if the α4 subunit is also phosphorylated by Ccnd1·Cdk *in vivo*, we immunoprecipitated Cdk substrates with an anti-phospho-Threonine-Proline (phosphoTP) antibody and the presence of phosphorylated α4 was assessed by western blot. We transiently transfected HEK-293T cells with α4, β3 subunits and Ccnd1 or Ccnd1^K112E^, a mutant of Ccnd1 that does not form an active complex with Cdk[40]. For the experiments where we co-expressed Ccnd1 or Ccnd1^K112E^, we used a variant that contains the CAAX motif of K-ras to maintain Ccnd1 attached to the cell membrane[10] to improve the phosphorylation of membrane targets. In the phosphoTP IP, we observed an increase in the phosphorylated α4 subunit in the presence of Ccnd1-CAAX relative to the presence of Ccnd1^K112E^-CAAX (Fig 2G and 2H). Moreover, when we used the non-phosphorylatable α4 mutant (α4^T423AS431A^: AA), we did not observe higher levels of phosphorylated α4 in the presence of Ccnd1 (Fig 2G and 2H). This data suggests that Ccnd1·Cdk4 can phosphorylate α4 *in vivo* at least at T423.

### Ccnd1·Cdk4 promotes the activation of α4-containing GABA_A_ receptors

Because Ccnd1·Cdk4 phosphorylates the α4 subunit in the intracellular loop, we reasoned that it may also affect the performance of α4-containing GABA_A_Rs. To study the electrophysiology of GABA_A_Rs we carried out whole cell patch-clamp recordings of HEK293 tsA201 cells transiently expressing α4 and β3 subunits (Fig 3A), since functional GABA_A_ receptors can be formed in HEK-293T cells by co-expression of these subunits[39]. To measure the behavior of the α4β3 GABA_A_Rs, 1 μM GABA was applied for 5 seconds every 2 min in these cells (Fig 3A and S1). Receptor-mediated current amplitude decreased (rundown) over time (Fig 3A and S1A): after 12 minutes of recording, GABA-mediated current was 60 ± 5 % (n=7) of the initial response (Fig 3A and S1D). Co-expression of Ccnd1-CAAX with α4β3 receptors partially prevented their rundown in a significant manner (Fig 3A and S1B). At 12 minutes after the start of the experiment, the GABA-mediated current amplitude, in the presence of Ccnd1-CAAX, was 80 ± 9 % (n=5) of the initial GABA-mediated response (Fig 3A and S1D). To confirm that phosphorylation of T423 S431 in α4 is important for this effect, we co-transfected a phosphomimetic α4^T423ES431E^ (α4^EE^) mutant with β3 and observed that it completely obliterates the rundown of GABA_A_Rs (Fig 3A and S1C). Thus, our data suggest that Ccnd1 can enhance the efficacy of α4-containing GABA_A_Rs by modulating the activity of the receptor through the phosphorylation of α4.

**Figure 3.**
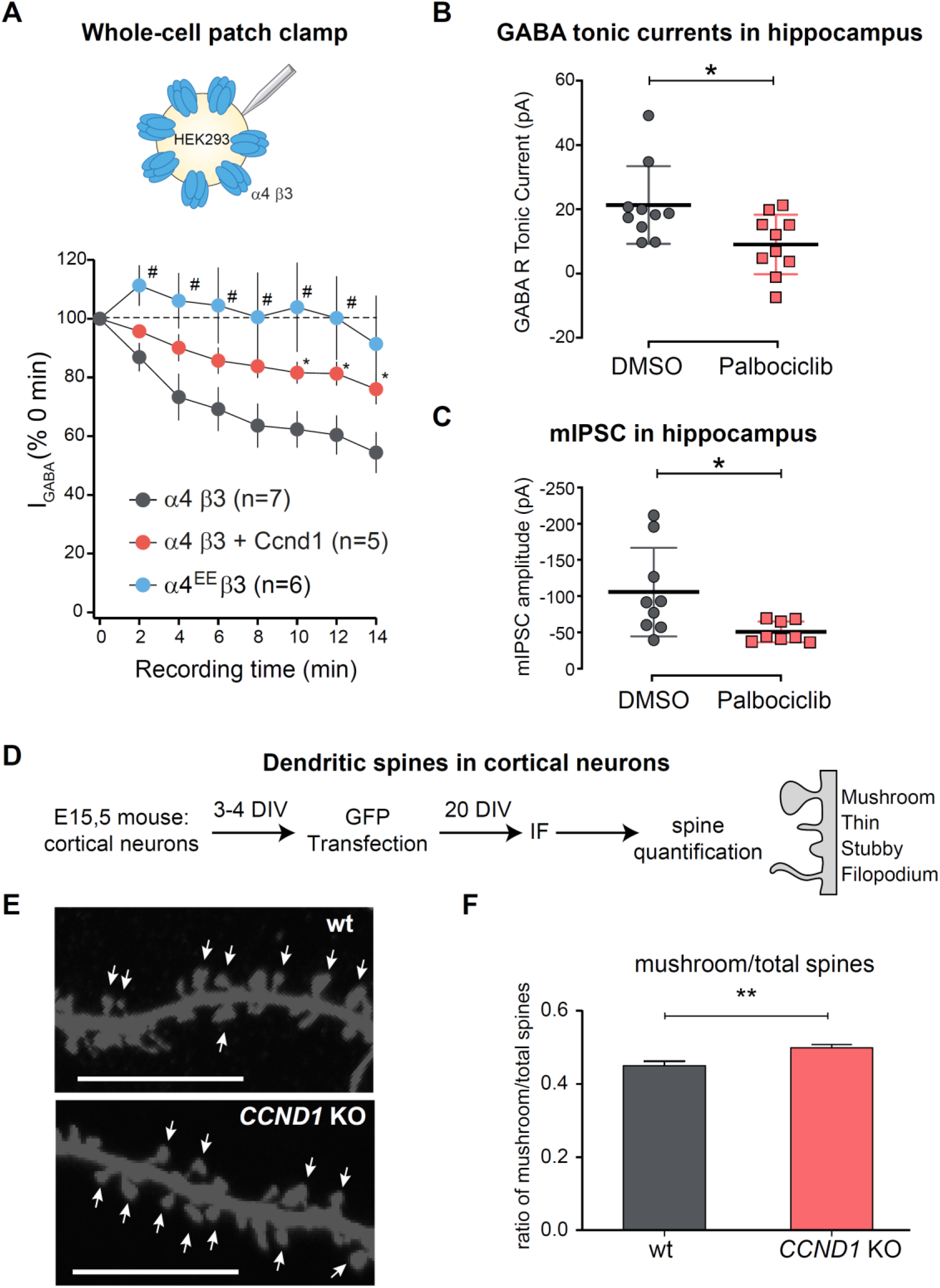
Ccnd1·Cdk4/6 modulates α4-containing GABA_A_ receptor activity. A) Ccnd1 or the phosphomimetic allele of α4 (α4^EE^) decreases GABA_A_ receptor rundown. Whole-cell patch clamp configuration for GABA currents recording in HEK-tsA201 cells transfected with α4 and β3 subunits of GABA_A_Rs. Currents were evoked by rapid application of 1 μM GABA and holding potential was held at −60mV. Peak current values at different times of GABA stimulations were analyzed in cells under 3 conditions: 1) α4 wt (α4β3, grey circles), 2) α4 wt co-transfected with Ccnd1-CAAX (α4β3+Ccnd1, red circles) and 3) α4^T423E S431E^ phosphomimetic allele (α4^EE^β3, blue circles). *p<0.05, α4β3 vs. α4β3+Ccnd1; #p<0.05 α4β3 vs. α4^EE^β3; Unpaired student’s t-Test. **B and C) Cdk4/6 inhibition by Palbociclib decreases GABA tonic currents and mIPSC amplitude in rat hippocampal slices**. GABA currents recordings in CA1 neurons: hippocampal slices from newborn rats were cultured in vitro, treated for 2 h with DMSO or Palbociclib 2.5 μM, and tonic (extrasynaptic) or mIPSCs (synaptic currents) were measured. B) GABA inhibitory postsynaptic currents were evoked by stimulation of Schaffer collaterals and the tonic GABA_A_ receptor-mediated current was measured as the outward shift in holding current following application of the GABA_A_R antagonist picrotoxin. CA1 pyramidal neurons were recorded under whole-cell voltage-clamp at -60 mV. Quantification of the shift in holding current for DMSO- and Palbociclib-treated cells (n=10), represented in a scatter dot plot showing mean values ± S.E.M (*p<0.05, Wilcoxon-Mann-Whitney test). The shift in holding currents, corresponding to tonic currents, is reduced in Palbociclib-treated slices compared to vehicle. C) To measure miniature inhibitory postsynaptic currents (mIPSC), 1 μM tetrodotoxin was added to block action potentials. Scatter dot plot showing mIPSC amplitude quantifications of DMSO-(n=9) and Palbociclib-treated cells (n=8). Bars show mean values ± S.E.M. *p<0.05, Wilcoxon-Mann-Whitney test. **D-F) CCND1 KO increases the maturity of dendritic spines in cortical neurons**. D) Schematic representation of the experimental design for dendritic spine analysis of cortical neurons at 20 DIV. Cultured cortical neurons from WT and CCND1 KO mice were transfected at 3 DIV with GFP and fixed at 20 DIV. Changes in spine density and morphology were assessed by analyzing GFP-positive neurons with NeuronJ (see Materials and Methods). E) Confocal images of representative neurites from WT and CCND1 KO neurons are shown. Scale bar: 10 μm. F) Quantification of the number of mushroom spines relative to the number of total spines from WT (n= 59 neurons) and CCND1 KO (n= 62) neurons (4 mice/group). Bars show mean values ± S.E.M. **p<0.01, Unpaired student’s t-test.

The α4 subunit is a component of extra-synaptic, high-affinity GABA_A_Rs, which mediate tonic inhibition. To assess the importance of Ccnd·Cdk4/6 on GABA_A_Rs in a physiological context, we measured tonic currents in hippocampal slices of newborn rats. Organotypic hippocampal slice cultures were treated with DMSO (vehicle) or the Cdk4/6 specific inhibitor Palbociclib at 2.5 μM for 2 h (Fig 3B and S2A). GABA inhibitory post synaptic currents were evoked with single voltage pulses in Schaffer collaterals, and tonic currents were recorded from CA1 pyramidal neurons under whole-cell voltage-clamp configuration. The application of the antagonist picrotoxin (100 μM) causes a decrease in the operative extra-synaptic GABA_A_ channels, which can be measured as an outward shift of the holding current[41] (Fig S2B). This shift corresponds to the GABA_A_R tonic currents (Fig 3B). While the holding current in DMSO-treated cells was shifted by 21 ± 4 pA, the shift in Palbociclib-treated slices was only 9 ± 3 pA (Fig 3B). Therefore, Ccnd·Cdk4/6 inhibition by acute Palbociclib application causes a significant decrease in the GABA_A_R tonic current.

Given that the α4 subunit can also form synaptic receptors when clustered together with βxγ2 subunits[42], we tested whether the acute Cdk4/6 inhibition may in turn affect GABA_A_ receptor synaptic function. To this end, we measured miniature inhibitory postsynaptic currents (mIPSC) in organotypic cultures treated with 2.5 μM Palbociclib for 2 h, in the presence of the action potential blocker tetrodotoxin (1 μM) (Fig S2A). We observed that the amplitude of the spontaneous miniature responses was reduced after Ccnd·Cdk4/6 inhibitor treatment as compared to the control (DMSO) treatment (-106 ± 20 pA for DMSO vs -51 ± 5 pA for Palbociclib; Fig 3C and S2C-D). Putting all these electrophysiological experiments together, our results suggest that Ccnd1·Cdk4/6 enhances GABA_A_R activity in the hippocampus.

Since Ccnd1 regulates α4-containing GABA_A_ receptors, we asked if the absence of Ccnd1 could disrupt some α4-dependent functions of GABA_A_Rs *in vivo*. The α4 subunit of GABA_A_Rs promotes synaptic pruning (the removal of dendritic spines) in the hippocampus of adolescent mice[27],[28]. To assess whether Ccnd1 could affect normal dendritic spine development, we analyzed the number and morphology of dendritic spines in cortical neurons from WT and *CCND1* KO mice. Cultured neurons were transfected with GFP at 3 DIV, and dendritic spines were analyzed at 20 DIV, when spines are completely developed (Fig 3D-F and S3). The density of dendritic spines did not depend on Ccnd1 expression (WT= 8.51 ± 0.16 vs *CCND1* KO= 8.28 ± 0.18 spines/10 μm, non-significant, Fig S3B). However, the ratio of the mushroom-shape (more mature) spines relative to total spines was significantly increased in *CCND1* KO neurons compared with neurons from WT animals (0.45 ± 0.01 vs 0.50 ± 0.01, p<0.01, Fig 3F). Overall, our data suggest that Ccnd1·Cdk4/6 enhances α4-dependent GABA_A_R activity, which may have consequences for normal dendritic spine development.

### Ccnd1 increases the surface levels of the α4 subunit of GABA_A_ receptors

Phosphorylation of several subunits of GABA_A_ receptors can affect their abundance at the cell surface, where they are active[43]. To analyze whether phosphorylation by Ccnd1·Cdk4 may affect GABA_A_R localization, we studied the surface levels of the α4 subunit when co-expressed heterologously with Ccnd1 in HEK-293T cells. For this, we transiently expressed α4 and β3 in these cells and assessed the surface levels of α4 by biotin labeling of surface proteins (Fig 4A). We used the transferrin receptor (TFRC) protein levels as a surface protein control and quantified the levels of α4 at the surface relative to the levels of α4 in the total cell extract (Fig 4B). We observed a statistically significant increase in the surface levels of α4 (148 ± 20%, P=0.04, Fig 4A and B) only when Ccnd1-CAAX was also co-expressed. This increase was abolished in the presence of Ccnd1^K112E^-CAAX, as well as when the non-phosphorylatable mutant α4^T423AS431A^ was used (98 ± 29%, P=0.93 and 107 ± 24%, P=0.82, respectively), indicating that the observed effect depends on T423/S431 phosphorylation by Ccnd1·Cdk4. These results suggest that Ccnd1 increases the surface levels of the α4 subunit of GABA_A_R in a kinase-dependent manner.

**Figure 4.**
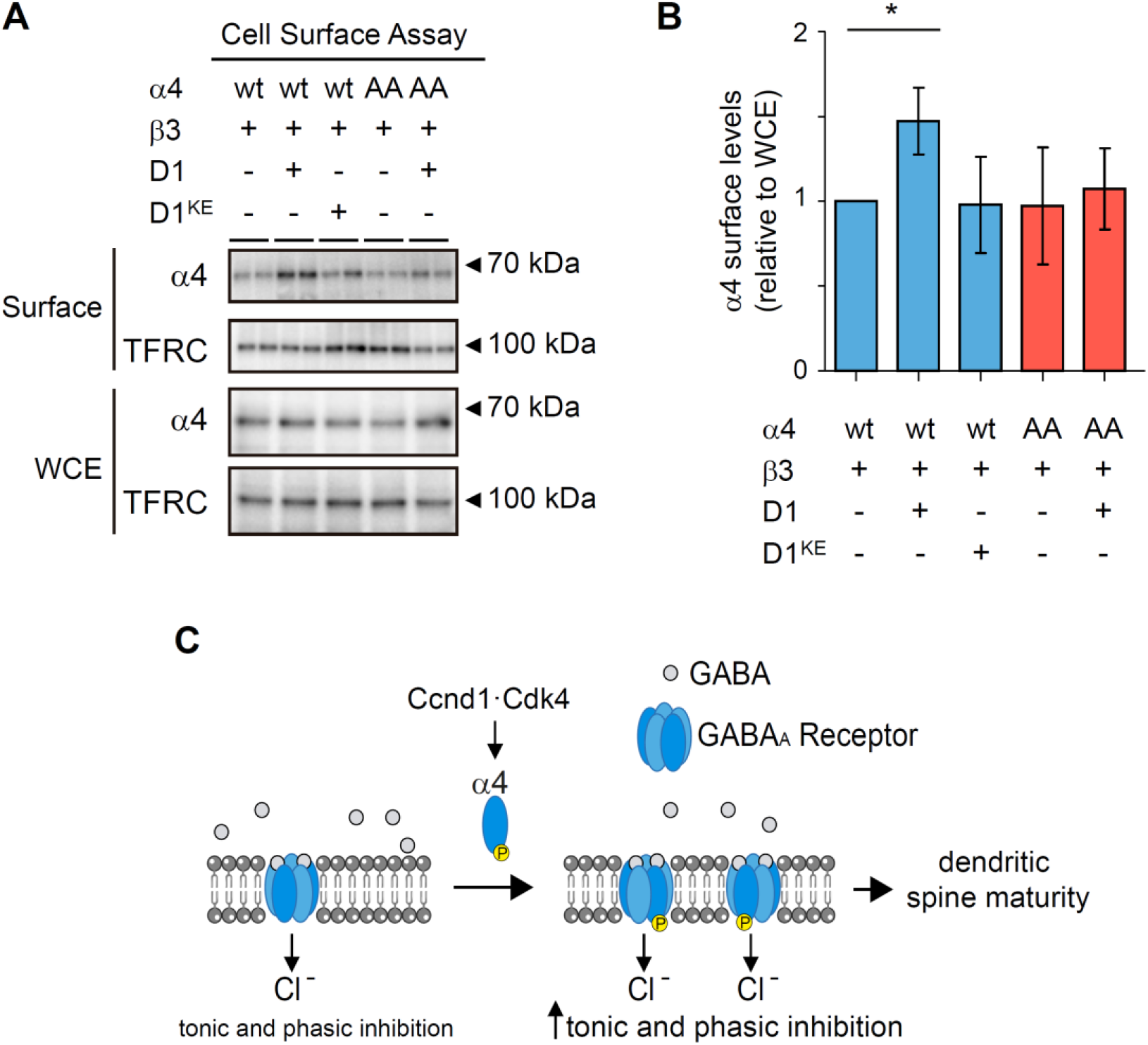
Ccnd1 increases surface levels of α4 subunit. A) HEK-293T cells were transfected with β3 and α4-FLAG or the nonphosphorylatable mutant (α4^T42A3S431A^-FLAG: AA). Either HA-Ccnd1-CAAX (D1) or HA-Ccnd1^K112E^-CAAX (D1^KE^) or a control vector was co-transfected. A biotinylation assay was used to purify cell surface proteins. TFRC is used to normalize α4 levels. Representative blots with whole cell extracts and surface proteins are shown. Antibodies used for protein detection are: polyclonal rabbit anti-FLAG to detect α4-FLAG; TFRC, monoclonal mouse anti-TFRC. B) Quantification of the experiment in A as a surface to WCE ratio. Bars show mean values ± S.E.M (n=4 independent experiments). *p<0.05 (Unpaired student’s t-test). C) Schematic model of the effects of Ccnd1·Cdk4 on α4-containing GABA_A_ receptors: Ccnd1·Cdk4 phosphorylates the α4 subunit, promoting its surface localization and activity, thus increasing synaptic (phasic) and extrasynaptic (tonic) inhibition and altering the maturity of dendritic spines.

## DISCUSSION

In neurons, Ccnd1·Cdk4 has been involved in promoting proliferation versus differentiation of stem cells during cortical development[13] and in the adult hippocampus[44]. However, the role of Ccnd1 in the regulation of neuronal signaling has never been investigated before. Interestingly, *CCND1* KO mice show an abnormal limb-clasping reflex[12], being this alteration an early symptom of neurological defects associated with hippocampal neuron malfunctioning[45]. In this work, we have unveiled a novel role of Ccnd1 in the stimulation of GABA_A_ receptor function (Fig 4C). Ccnd1 interacts with the α4 subunit of GABA_A_R and Ccnd1·Cdk4 complex phosphorylates T423 and S431 in the intracellular loop of α4. This phosphorylation increases the surface levels of α4 subunit and the efficacy of GABA_A_R activity in whole cell patch-clamp recordings. Furthermore, Ccnd·Cdk4/6 inhibition decreases tonic currents and the amplitude of mIPSCs in the hippocampus of newborn rats. Accordingly, the proportion of mature dendritic spines is increased in cortical neurons from *CCND1* KO mice. Therefore, we propose that Ccnd1·Cdk4 mediated phosphorylation is an important physiological regulator of GABA_A_R-mediated inhibition.

Ccnd1 has been found in the cytoplasm of postmitotic neurons[16]. As nuclear localization of Ccnd1 in these cells promotes apoptosis, its cytoplasmic localization has been suggested as a Ccnd1 sequestration mechanism to avoid cell death. However, Ccnd1·Cdk4 has cytoplasmic functions interacting with membrane-associated proteins such as Filamin A, RalGTPases and Paxillin[6],[8],[9],[46]. It is then possible that Ccnd1·Cdk4 plays an active role in the cytoplasm of post-mitotic neurons. Supporting this, the mRNA of Ccnd1 has also been found in the synaptic neuropil in the hippocampus[47], suggesting local translation of Ccnd1 in the dendrites. Also, Cdk4 has been localized in the cytoplasm in adolescent mouse hippocampal neurons[20]. According with a hippocampal cytoplasmic expression of Ccnd1, here we have detected co-immunoprecipitation of Ccnd1 and α4 in samples from mouse hippocampus where this subunit of the GABA_A_R is abundantly expressed[23], providing evidences for a cytoplasmic role of Ccnd1 in neurons.

In this work, we have proved that Ccnd1·Cdk4 phosphorylates α4 at T423 and S431 in the intracellular loop (ICL) *in vitro*. Moreover, in transiently transfected HEK-293T cells, we have detected Ccnd1-specific phosphorylation of α4 *in vivo*. Rundown of GABA_A_ receptors involves phosphorylation-dephosphorylation processes[48],[49],[50]. The ICL between TM3 and TM4 of several subunits of GABA_A_Rs is regulated by phosphorylation, affecting the interaction of GABA_A_ receptors with other proteins, the stability and the surface localization of the subunits, or the activity of the GABA_A_ receptors[21]. For instance, PKC phosphorylates the α4 subunit in the intracellular loop (S443) increasing its stability, membrane insertion and the efficacy of GABA_A_Rs[39],[51]. Consistent with these previous data, we have determined by whole-cell patch-clamp recordings that the expression of Ccnd1 or the phosphomimetic allele of α4 (α4^T423ES431E^) caused a decrease in the rundown of α4β3 containing GABA_A_ receptors in HEK-293 cells. We propose that Ccnd1·Cdk4 enhances the activity of α4-containing GABA_A_ receptors by promoting the surface accumulation of α4 through its phosphorylation. On the one hand, Ccnd1 but not the inactive allele Ccnd1^K112E^ is able to promote surface accumulation of α4. Moreover, Ccnd1 does not increase surface levels of the non-phosphorylatable mutant of α4 (α4^T423AS431A^).

The adjustment of the surface levels of GABA_A_R subunits takes place through the regulation of export and internalization processes. The ICL of β3 subunit is involved in internalization of the receptors by clathrin. There is an atypical site for interaction with clathrin adaptor protein 2 (AP2) (KTHLRRRSSQLK), that contains the phosphorylation site for PKA, PKC and CamKII: it only binds AP2 when dephosphorylated[52]. Such a mechanism could also be happening for α4 subunit, that contains a similar motif (NMRKRTNAL) in the intracellular loop (aa 385-393). In fact, α4 has been described to interact directly with clathrin μ2 by immunoprecipitation[53]. Although T423 and S431 are not found inside the motif, phosphorylation of the ICL and/or the presence of Ccnd1 could be affecting the binding of α4 to clathrin. Another possibility is that the trafficking from the ER to the cytoplasmic membrane is affected. In the α4 and δ subunits there are dibasic or multibasic motifs in homologous positions that reduce export from the ER and increase transport back from the Golgi apparatus to the ER[54]. The mutation of dibasic motif to Ala in α4 enhances surface expression of the receptors. Again, phosphorylation of ICL or interaction with Ccnd1 could affect the association of dibasic sites with trafficking proteins. Consistently with a role of Ccnd1·Cdk4/6 on GABAergic signaling, we have shown that inhibition of Cdk4/6 in CA1 pyramidal neurons produces a decrease in tonic currents and the amplitude of mIPSCs. Even though α4 is mostly extrasynaptical, it can also form synaptic GABA_A_Rs when assembled with γ2. Concerning mIPSC amplitude, it is feasible that phosphorylation of α4 by Ccnd·Cdk4/6 complexes is still promoting its activity also when the α4 subunit is forming α4βxγ2 receptors at the synapse[55]. This would mean that α4 phosphorylation by Ccnd·Cdk4/6 has the same effect irrespective of whether the receptor localizes synaptically or extrasynaptically. By contrast, α4-containing synaptic and extrasynaptic receptors show different modulation by PKA and PKC phosphorylation. PKA activation decreases synaptic α4 expression while increases extrasynaptic α4. Activation of PKC instead promotes α4-expression at the synapse and has no effect extrasynaptically[56]. Dendritic spines are the postsynaptic sites where most excitatory synapsis occurs. Their change in number and morphology underlies synaptic plasticity, important for learning and memory processes. GABA, through GABA_A_Rs activation promotes spine shrinkage and elimination[57]. Specifically, GABA_A_Rs containing α4βδ subunits have been involved in synaptic pruning at the CA1 and CA3 regions in the hippocampus[27],[28]: the expression of the α4 subunit of GABA_A_Rs increases in the hippocampus during adolescence[28] and the density of dendritic spines decreases post-pubertal, while in the *GABRA4* KO mice this pruning does not occur. Therefore, the density of dendritic spines in the hippocampus of adult mice is increased in the *GABRA4* KO mice[37]. The effect of α4 on synaptic pruning is mainly observed in mushroom- and stubby-shaped dendritic spines[27],[28]. According to a role of Ccnd1 in promoting α4-containing GABA_A_Rs activity, we observed an increase in the proportion of mushroom-type spines in *CCND1* KO neurons. Mushroom spines are considered to be one of the mature types of spines, involved in memory[58]. In this sense, the increase in α4βδ receptors at puberty impairs learning[25] and the *GABRA4* KO mouse has increased spatial memory[37]. In the same direction, increased expression of Ccnd1 was found in CA1 region of aged learning-impaired rats[59] and the expression of Ccnd1 in the hippocampus of rats triggers deficits in spatial working memory[60]. Moreover, *GABRA4* KO mice show an autistic-like behavior, and autism has been associated with an increased number of spines[61]. The effect of Ccnd1 on GABA signaling and dendritic spines opens the possibility of exploring Ccnd1 functions in diseases related to the imbalance between inhibitory/excitatory signaling and dendritic spine abnormalities such as autism spectrum disorders and epilepsy.

## MATERIALS and METHODS

### Cell culture

Mycoplasma-free HEK293T cells were obtained from the American Type Culture Collection. Cells were maintained at 37ºC in a 5% CO_2_ incubator, and grown in Dulbecco’s modified Eagle’s medium (DMEM) supplemented with 10% Fetal Bovine Serum (FBS), 10 U/mL penicillin/streptomycin (P/S) and 4 mM glutamine. Mycoplasma detection tests were performed in the Cell culture service (SCT-CC). Transient transfection of vectors was performed with Lipofectamine 2000 (Invitrogen) according to manufacturer’s instructions.

### Expression vectors

Human CCND1 was fused to three copies of the Flag or HA or one of Red monomer epitope under the CMV promoter in pcDNA3. Ccnd1-CAAX construct is a fusion of the 3’ end of the K-Ras ORF containing the CAAX motif (GGC TGT GTG AAA ATT AAA AAA TGC ATT ATA ATG TAA) to the 3’ end of the CCND1 ORF[10]. Mouse GABRA4 (IMAGE ID 6828002, Source BioScience) and Human GABRB3 (IMAGE ID 3871111, Source BioScience) were used to obtain N-terminal GST fusions in pGEX-KG (Clontech) or N-terminal FLAG fusions in pJEN1 (pcDNA3 derived, from Dr. Egea) or cloned into FCIV1 lentiviral vector (from Dr. Encinas). Standard PCR-mediated site-directed mutagenesis was used to obtain the non-phosphorylatable and phosphomimetic mutants of GABRA4 using the following primers: T423A and S431A: CGCCAAATCCATTCAGCAGGGC and CCGAAGCTAAGTGAGCCTTAGGCGCGGCTTCAGAAGACTCCTGGAC; S456A and S458A: CGGCCCCCGCTCCTCATGGCACATTGCGGC and CGGCAGATGAAAGACCTCTGGCTGM; T500A: CGCCTCCTCCCCCTGCTCCAC and CCGCTGACACATTCCCAGCAGC; and T423E and S431E: GCGAACCAAATCCATTCAGCAGGGC and TAGCTAAGTGAGCCTTAGGCTCGGCTTCAGAAGACTCCTGGAC.

### Immunoprecipitation assays

For co-immunoprecipitation experiments, HEK293T cells were harvested 24 h after transfection. The cells were resuspended in RIPA buffer (50mM Tris pH7.4, 150mM NaCl, 1% NP-40, 0.1% sodium deoxycholate, 0.1% SDS, 5mM EDTA, protease and phosphatase inhibitors), rocked 1 h at 4ºC, and spun for 5 min at 600 g. The supernatants were incubated with protein G magnetic beads (Dynabeads, Invitrogen) for 30’ at 4ºC. After the preclearing, the samples were incubated with 2 μg of αFLAG M2 antibody (Sigma) overnight and precipitated with protein G dynabeads. IP of anti-phosphoThreonineProline antibody (#9391, Cell Signaling) was performed similarly, but using protein A dynabeads. The beads were collected and washed three times with 1 mL of cold RIPA buffer, and bound proteins were separated by SDS-PAGE gels and visualized by western blots.

IP of endogenous Ccnd1 was carried out in hippocampus of adult female mice with a polyclonal antibody (#06–137, Upstate). Briefly, cleared cell extracts in RIPA buffer (with protease and phosphatase inhibitors) were immunoprecipitated with Protein A linked to magnetic beads (Invitrogen). An anti-FLAG (#F7425, Sigma) was used as a mock control. Washes were performed with RIPA buffer without SDS.

### Immunoblotting

For immunoblot, protein samples were resolved by SDS-PAGE, transferred to PVDF membranes (Millipore), and incubated with primary antibodies: anti-Ccnd1 (monoclonal DCS-6, BD biosciences, 1:500), anti-Gabra4 (polyclonal, AB5459, Merck, 1:1000, and polyclonal 12979-1-AP, Proteintech, 1:500), anti-Flag (polyclonal, 11508721, Invitrogen, 1:1000), anti-HA (rat monoclonal 3F10, Roche #11867431001, 1:5000), anti-TFR (monoclonal H68.4, Invitrogen #13–6800, 1:1,000) and anti-GST (goat polyclonal, Amersham #27–4577, 1:2000). Appropriate peroxidase-linked secondary antibodies (GE Healthcare UK Ltd) were detected using the chemiluminescent HRP substrate Immobilon Western (Millipore). Chemiluminescence was recorded with a ChemiDoc-MP imaging system (BioRad).

### Production and purification of recombinant proteins

All of the GST fusions were expressed in *E. coli* BL21 (DE3) by adding 1 mM isopropyl-β-D-thiogalactopyranoside (IPTG) to LB broth cultures at a cell density of 0.3 A_600_ and subsequent incubation for 4 h at 30 °C. The proteins were purified using glutathione-Sepharose 4B beads as directed by the supplier (Amersham Biosciences) in 500 μl of lysis buffer containing 50mM HEPES, pH 7.5, 150 mM NaCl, 1 mM EDTA, 0.5% Triton X-100, 10% glycerol, 1 mM DTT, and protease and phosphatase inhibitors. The concentration and purity of substrates were estimated by comparison to protein standards stained with Coomassie Brilliant Blue.

For *in vitro* transcription and translation, FLAG-Ccnd1 was amplified by PCR (forward primer: CGCGCTAATACGACTCACTATAGGGAGACCCAAGCCCATGGGATCAC; reverse primer: TTTTTTTTTTTTTTTTTTTTTTTTTTTTTGGCTGATCAGCGAGCTCTAG), transcribed *in vitro* with the T7 RNA polymerase (New England Biolabs), and *in vitro* translated with a Rabbit Reticulocyte Lysate system (Promega).

### GST pull-down assay

For the GST pull-down assay, 400 ng of GST or GST-α4 or GST-Cter-α4 purified from *E. coli* immobilized on Glutathione-Sepharose 4B beads were incubated with Flag-Ccnd1 in binding buffer (20 mM HEPES-KOH, pH 7.5, 150 mM KCl, 5 mM MgCl_2_, 0.5 mM EDTA, 0.1% NP-40, 1 mM DTT, 1 mM PMSF, 10% glycerol, protease and phosphatase inhibitors) for 30 min at room temperature. After four washes with the same buffer, the samples were analyzed by SDS-PAGE.

### Kinase assay

Briefly, 0.2 mg substrate (either GST-α4, GST-Cter-α4 or GST-β3) was mixed with 1.5 μL of active Ccnd1-Cdk4 complex purified from baculovirus (Sigma C0620), 10 mM ATP, 7 mCi of γ-^32^P-ATP (PerkinElmer, 3000 Ci/mmol), and either DMSO or 2 μM of the Cdk4/6-inhibitor Palbociclib (Selleckchem, S1116) in 20 μL of kinase buffer (50 mM Tris–HCl pH 7.5, 10 mM MgCl_2_, 0.5 mM DTT, 1 mM EGTA and 2.5 mM b-glycerophosphate). This mixture was incubated for 20 min at 30 ºC, then boiled in 2x Laemmli buffer, and separated by electrophoresis. Phosphorylated proteins were visualized by autoradiography of the dried slab gels.

### MS analysis of phosphopeptides

The C terminus of α4 fused to GST was used in an *in vitro* kinase assay with Ccnd1·Cdk4 in the presence of ATP or in the absence of ATP as a control. Samples were subsequently subjected to SDS-PAGE. The gels were stained with Coomassie Brilliant Blue G-250 colloidal (EZBlue Gel Staining Reagent, Sigma). After washing with water, protein bands of interest were prepared and submitted to Fundació Institut d’Investigació Biomèdica de Bellvitge (IDIBELL) Proteomics Service (Barcelona) for analysis of the Chymotryptic peptide molecular masses by liquid chromatography-mass spectrometry. Gel slices were manually cut and proteomic service suggested for each band-assay to recover three slices (low 1, middle 2, and high 3 mobility) as phosphorylation could alter band mobility. Briefly, gel bands were washed with water, ammonium bicarbonate (50mM) and 50% acetonitrile. Next, samples were reduced by incubation with DTT (10 mM) at 60 ºC for 45 min and alkylated with iodoacetamide (50mM) for 30 min, in the dark. Finally, proteins were digested with chymotrypsin (5 ng/μL) at 25ºC overnight (Trypsin gold, Promega). Digestion was stopped by addition of 5% formic acid and peptides extracted twice with 70% acetonitrile and 5% formic acid (10 min sonication). Peptide extracts were evaporated to dryness, resuspended with 2% acetonitrile 0.1% formic acid and analyzed by nano-HPLC-MSMS.

### Surface biotinylation

For surface biotinylation (all steps performed on ice) HEK-293T cells were washed twice with PBS and incubated with 1 mg/mL EZ-link sulfo-NHS-SS-biotin (Pierce), a nonpermeable biotin, in PBS for 15 min. Following surface labeling, nonconjugated biotin was quenched by washing twice with TBS. Cells were lysed in RIPA buffer (50 mM Tris, pH7.4, 150 mM NaCl, 1% NP-40, 0.1% SDS, 5 mM EDTA, with phosphatase and protease inhibitors), and protein concentrations were determined using a DC protein assay (Bio-Rad). Equal amounts of protein (usually 1 mg in 500 μL) were incubated overnight with 20 μL NeutrAvidin-coated beads (Pierce) at 4°C with constant rotation. The next day, beads were washed three times with RIPA buffer and eluted samples were processed for immunoblotting.

### Whole-cell currents measurements in tsA201 cells

Electrophysiological experiments were done with mycoplasma-free tsA201 cells (kindly provided by Prof. F. Ciruela, Universitat de Barcelona) and purchased from the American Type Culture Collection (ATCC, CRL-3216, RRID: CVCL_0063). Mycoplasma detection tests were performed with PlasmoTest (Invivogen, code: rep-pt1). Cells were transiently co-transfected with 1 μg total cDNA using polyethylenimine (PEI) transfection reagent (1 mg/mL) in a 3:1 ratio (PEI:DNA). In all transfections the DNA ratio used was 1:1:2 (plasmid A: plasmid B: plasmid C), where A is α4 or α4^T423ES431E^, B is β3 and C is a control vector or Ccnd1-CAAX codifying plasmid. Cells were re-plated on poly-D-lysine coated glass coverslips to allow optimal density with isolated cells. All experiments were performed 48 h after transfection.

For the measurements of GABA-evoked whole-cell currents in tsA201, cells were visualized with an inverted microscope (AxioVert A.1, Carl Zeiss) and maintained in extracellular flowing solution at a rate of 60 mL/h. The extracellular solution contained (in mM): 140 NaCl, 5 KCl, 10 HEPES, 11 glucose, 2.5 CaCl_2_, 1.2 MgCl_2_ (pH = 7.4 with NaOH; osmolarity 305 mOsm/Kg adjusted with sorbitol). Electrodes were fabricated from borosilicate glass (1.50 mm O.D., 1.16 I.D., Harvard Apparatus) pulled with a P-97 horizontal puller (Sutter Instruments) and polished with a forge (MF-830, Narishige) to a final resistance of 2–5 MΩ. The intracellular solution contained (in mM): 140 KCl, 2 MgCl_2_, 2 CaCl_2_, 10 HEPES, 1.1 EGTA, 2 Mg ATP (pH = 7.2 with KOH; osmolarity 295 mOsm/Kg adjusted with sorbitol).

Macroscopic GABA_A_-mediated currents were recorded at room temperature (22-25°C) in the whole-cell configuration from cells positive for Venus (α4 in FCIV1) and mCherry (Ccnd1-CAAX in pcDNA3 Red monomer) fluorescence at a holding membrane potential of -60 mV. Currents were recorded with Axopatch 200B amplifier, filtered at 2 kHz and digitized at 5 kHz using Digidata 1440A interface with pClamp 10 software (Molecular Devices Corporation). Series resistance was typically 5–20 MΩ, and was monitored at the beginning and at the end of the experiment. Cells that showed a change in series resistance greater than 20% were rejected. Rapid application of the agonist GABA (1-2 ms exchange) was applied by switching between a continuously flowing control solution (extracellular solution diluted by 5%) and a GABA-containing solution (1 μM GABA and 2.5mg/mL of sucrose diluted in extracellular solution). Solution switching was achieved by piezoelectric translation of a theta-barrel application tool made from borosilicate glass (1.5 mm O.D.; Sutter Instruments) mounted on a piezoelectric translator (P-601.30; Physik Instrumente). Agonist was applied at 2 minutes interval and the magnitude of the peak current was measured (I_GABA_). Electrophysiological recordings were analyzed using IGOR Pro (Wavemetrics Inc.) with NeuroMatic (Jason Rothman, UCL).

### Preparation of organotypic hippocampal slice cultures

Organotypic hippocampal slice cultures were prepared as previously described[62]. Briefly, whole brain of newborn rats (postnatal days 5-7) were dissected and placed in ice-cold Ca^2+^-free dissection solution (10 mM D-glucose, 4 mM KCl, 26 mM NaHCO_3_, 8% sucrose, 5 mM MgCl_2_, 1:1000 Phenol Red) saturated with 5% CO_2_ / 95% O_2_, enabling the brain to chill for 1 min. Hippocampi were then isolated with the help of a magnifying glass, cut in slices (400 μm) in the same solution using a tissue chopper (Stoelting), and maintained at 35.5 °C in culture on permeable membranes in a medium containing 20% horse serum, 1 mM CaCl_2_, 2 mM MgSO_4_, 1 mg/L insulin, 0.0012% ascorbic acid, 30 mM Hepes, 13 mM D-glucose, and 5.2 mM NaHCO_3_. Culture medium was replaced with fresh one every 2-3 days. The slices were used at 4-8 days *in vitro*.

### Whole-cell recordings of GABA_A_ receptor-mediated currents

Prior to the electrophysiological recordings, slices were treated for 2 hours with either DMSO (control solution) or 2.5 μM Palbociclib (Cdk4/6 inhibitor) (Selleckchem). After that, slices were placed in a chamber and perfused with artificial cerebrospinal fluid [aCSF: 119 mM NaCl, 2.5 mM KCl, 1 mM NaH_2_PO_4_, 26 mM NaHCO_3_, 11 mM glucose, 1.2 mM MgCl_2_, 2.5 mM CaCl_2_; osmolarity adjusted to 290 mOsm; pH7.5] supplemented with 100 μM AP5 (NMDA receptor antagonist), 10 μM CNQX (AMPA receptor antagonist), 1 μM strychnine (glycine receptor antagonist), 4 μM 2-chloroadenosine (to reduce neurotransmitter release and stabilize synaptic transmission) and DMSO or Palbociclib as external solution gassed with 5% CO_2_ / 95% O_2_ at 29 °C in the electrophysiology set-up. Patch recording pipettes (4-6 MΩ) were pulled from thin-walled borosilicate capillary glass (World Precision Instruments [WPI], Sarasota, FL) on a P-2000 laser electrode puller (Sutter Instrument, San Rafael, CA) and filled with high chloride internal solution (178 mM CsCl, 10 mM Hepes, 2.5 mM MgCl_2_, 4 mM Na_2_ATP, 0.4 mM Na_3_GTP, 10 mM sodium phosphocreatine, 0.6 mM EGTA; pH adjusted to 7.2, osmolarity adjusted to 290 mOsm). GABA inhibitory postsynaptic currents were evoked with single voltage pulses (200 μs, <30 V) delivered through platinum-iridium bipolar electrodes (FHC, Bowdoin, ME, USA) placed on Schaffer collaterals. CA1 pyramidal neurons were recorded under whole-cell voltage-clamp at -60 mV using Multiclamp 700A amplifier (Molecular Devices, San Jose, CA, USA). At least 10 min of stable baseline synaptic responses and holding currents were recorded. The tonic GABA_A_ receptor-mediated current was measured as the outward shift in holding current following application of picrotoxin (100 μM). Synaptic responses were used to monitor effective inhibition of GABA_A_Rs by picrotoxin. To measure miniature inhibitory postsynaptic currents (mIPSC), 1 μM tetrodotoxin was added to the aCSF to block action potentials. Recordings were obtained with the gap-free mode of pClamp and mIPSCs were analyzed with the event detection of pCLAMP software (Molecular Devices).

### Cortical neuron culture

For primary culture of cortical neurons, cortexes were dissected from E15.5 *CCND1* KO or WT mice in ice-cold HBSS-MHPS (Hank’s Balanced Salt Solution, 10 mM MgSO_4_, 10 mM HEPES pH 7.2 and 10 U/mL P/S). After dissection, cortexes were incubated with papain solution (1 mg/mL papain in HBSS-MHPS) at 37ºC for 23 min. Enzymatic digestion was inhibited by washing the tissue 3 times with 10 mg/mL trypsin inhibitor (Fisher) in HBSS-MHPS. Additionally, one more wash was done with Neurobasal medium supplemented with 2% B27, 2 mM Glutamax and 1 mM sodium pyruvate (NB27, Fisher) at room temperature. The tissue was triturated by passing it through a flame-polished Pasteur pipette (<20 times). After the mechanical disaggregation, the supernatant containing mostly single cells was centrifuged at 650 rpm for 4 min. The pellet was resuspended in NB27 and the cells were counted in a Neubauer chamber. Cells were seeded at low density (25000 cells/cm^2^) with NB27 medium in poly-D-lysine (0.5 mg/mL) and laminin (5 μg/mL)-coated plates, in 1:1 conditioned medium. One third of the culture medium is replaced with fresh one every 3-4 days. Primary cortical neurons were transfected at 3 DIV using Lipofectamine 2000 (Invitrogen) following the manufacturer’s recommendations and the proportion 0.3 μg DNA: 1.2 μL Lipofectamine2000. The media was changed 4 hours after transfection and cells were fixed at 20 DIV.

### Genotyping of *CCND1* knockout mice

*CCND1* knockout mice[12] were obtained from Charles-River. Transgenic mice were maintained in a mixed background of C57BL/6 and SV129. Embryo’s tails were collected and warmed at 95 ºC NaOH 50 mM for 45 min. After neutralization with Tris 0.15 M pH8.5, the samples were then centrifuged for 1 minute at 10000 rpm. Supernatant containing genomic DNA was used for CCND1 genotyping by PCR. The following primers were used: a common primer CTGTCGGCGCAGTAGCAGAGAGCTACAGAC, a CCND1 primer CGCACAGGTCTCCTCCGTCTTGAGCATGGC and a neomycin primer CTAGTGAGACGTGCTACTTCCATTTGTCACG. PCR was performed in a Thermal Cycler (T100 Bio-Rad) and PCR products were analyzed in a 1.5 % agarose gel followed by Ethidium Bromide staining. The expected band pattern is 249 bp for WT, and 394 bp for KO alleles.

### Immunofluorescence and spine analysis

Cortical neurons were fixed using 4% PFA and 4% sucrose in PBS for 15 min at room temperature and then washed with PBS. Neurons were permeabilized for 5 min with 0.1% Triton X-100 in PBS and blocked with 3% BSA in PBS. Primary antibody α-GFP (A11120, ThermoFisher) was diluted 1:200 in 0.3% BSA in PBS. Proteins were detected by incubation with secondary antibody Alexa 488 rabbit anti-mouse (A11059, ThermoFisher). Images were acquired with an Olympus FV1000 confocal microscope using the following parameters: stacks of 10 slices were imaged every 0.37 μm, under a 60X objective. Laser power and PMT values were kept constant throughout images and conditions. Immunofluorescence quantification was performed using NeuronJ of ImageJ. Spine density and morphology analysis of WT and *CCND1* KO cortical neurons was performed on 3D reconstructions of confocal z series acquired using a 2x zoom. Spines were counted as mushroom-type if the spine head was wider than the spine neck. All analysis was done blind to the experimental condition.

### Statistical analyses

Different statistical analyses were used in this work. In every analysis, the minimum level of statistical significance was a p-value equal to 0.05. The significance level is represented in each graph as indicated. In order to evaluate possible differences between two experimental groups a t-test was performed. When there was a multiple comparison, one-way ANOVA with a post-hoc test (Tukey) was performed. In whole-cell recordings in ts201A cells, comparisons between groups were done using the parametric student’s t-test. For tonic currents comparison, Wilcoxon test was performed and in mIPSC amplitude, Mann-Whitney and Kolmogorov-Smirnov tests were done. Microsoft Excel and GraphPad Prism v5.0 (GraphPad) were used for statistical analysis and graphical representation.

### Ethical considerations

All procedures with animals follow the protocols approved by the Institutional Committee of Care and Use of Animals (Comitè Institucional de Cura i Ús d’Animals), and experiments were approved by the Ethics Committee of the University of Lleida (CEEA 03-02/19). Animals were housed and maintained in the animal facility of the University of Lleida with 12h:12h light/dark cycle and food/water available *ad libitum*. For electrophysiological experiments, all biosafety procedures and animal care protocols were approved by the bioethics committee from the Consejo Superior de Investigaciones Cientificas and were performed according to the guidelines set out in the European Community Council Directives (2010/63/EU, 86/609/EEC). All efforts were made to minimize the number of animals and their suffering.

## ACKNOWLEDGEMENTS

We are grateful to M. Encinas for plasmids. We thank Sònia Rius for technical assistance and members of the CYC lab for helpful considerations. The cell culture experiments were performed in the Cell Culture Scientific & technical Service from Universitat de Lleida. This work was funded by the Spanish Ministry of Innovation and Science MICINN (PID2019-104859GB-I00 to E.G. and PID2020-117651RB to J.A.E.) and by Generalitat de Catalunya (2017-SGR-569). M. Ventura Monserrat was supported by a predoctoral fellowship of University of Lleida.

## AUTHORS CONTRIBUTION

Neus Pedraza: conceptualization, supervision, validation, investigation, methodology, writing-original draft, writing – review & editing; Ma Ventura Monserrat: validation, investigation, methodology; Francisco Ferrezuelo: funding acquisition, data curation, writing – review & editing; Jordi Torres: funding acquisition, supervision, writing – review & editing; Neus Colomina: funding acquisition, methodology; David Soto: funding acquisition, investigation, validation, methodology; Fede Miguez: investigation, methodology; JA Esteban: supervision, validation, methodology, funding acquisition; Joaquim Egea: funding acquisition, conceptualization, methodology; Eloi Garí: conceptualization, supervision, funding acquisition, investigation, writing – review & editing.

---

**S1 Table.**
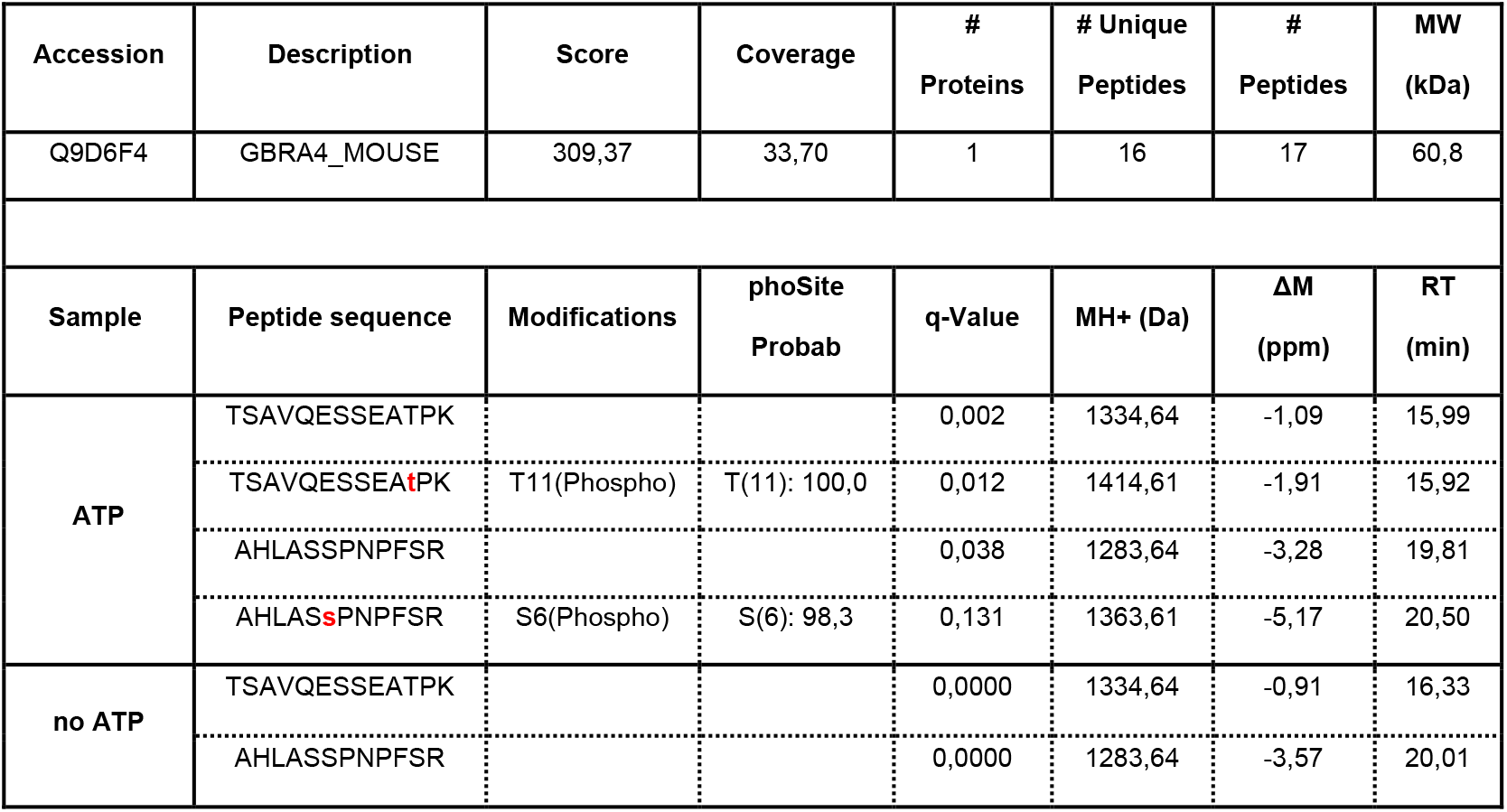
The α4 subunit of GABA_A_ receptors is phosphorylated at T423 and S431 by Ccnd1·Cdk4. The GST-Ct-α4 wild type was used in an in vitro kinase assay with Ccnd1·Cdk4, in the presence or absence of ATP, and the Chymotryptic peptide molecular masses were analyzed by liquid chromatography-mass spectrometry. MH+: proton adduct; ΔM: mass difference; RT: retention time; q-value: adjusted p-value. Both T423 and S431 of α4 subunit (highlighted in red) were found to be phosphorylated by Ccnd1·Cdk4.

**S1 Figure.**
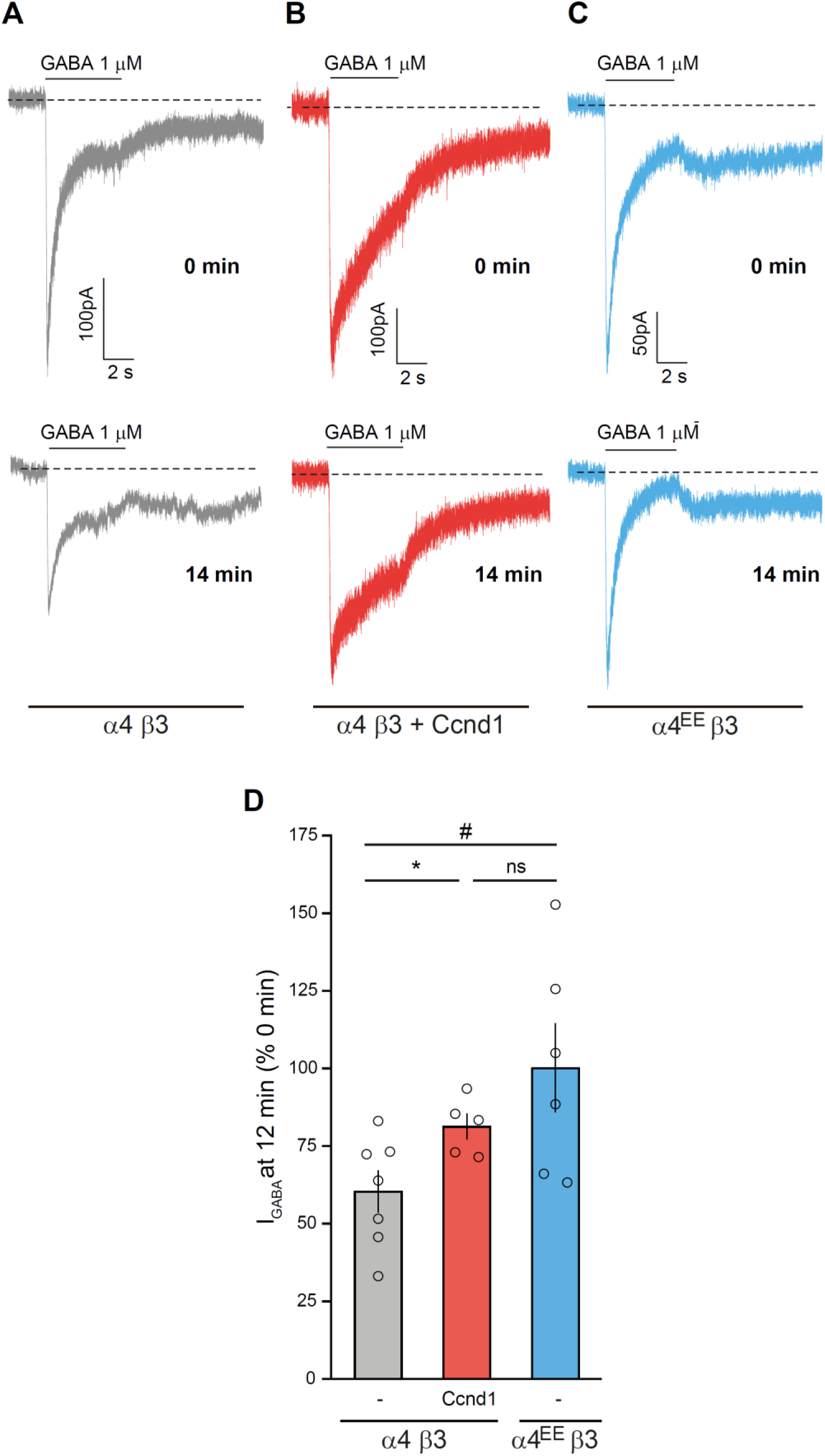
Ccnd1 or the phosphomimetic allele of α4 (α4^EE^) decreases GABA_A_ receptor rundown. A) Representative recording of whole-cell currents evoked by rapid application of 1 μM GABA (solid line above the trace) to HEK-tsA201 cells expressing wild-type α4β3 GABA_A_Rs. The GABA-evoked current at the beginning of the recording (time 0 min) and the response after 14 minutes (time 14 min) are shown. Dashed line denotes the zero current. Holding potential was held at −60mV. B) Same as A, but from cells co-expressing wild-type α4β3 GABA_A_Rs plus Ccnd1-CAAX. Note that the rundown of the peak current at 14 minutes is smaller than in condition A. C) Same as A, but from cells co-expressing phosphomimetic (T423E S431E) GABA_A_ receptor α4^EE^β3. D) Bar graph showing the mean ± S.E.M. of the relative current at t=12 minutes compared with current at time=0 for cells expressing wild type α4β3 GABA_A_Rs (n=7), α4β3 GABA_A_Rs plus Ccnd1-CAAX (n=5) or for α4^EE^β3 GABA_A_Rs (n=6). Open circles denote single experimental values. *p<0.05, α4β3 vs. α4β3+Ccnd1-CAAX; #p<0.05 α4β3 vs. α4^EE^β3; Unpaired student’s t-Test.

**S2 Figure.**
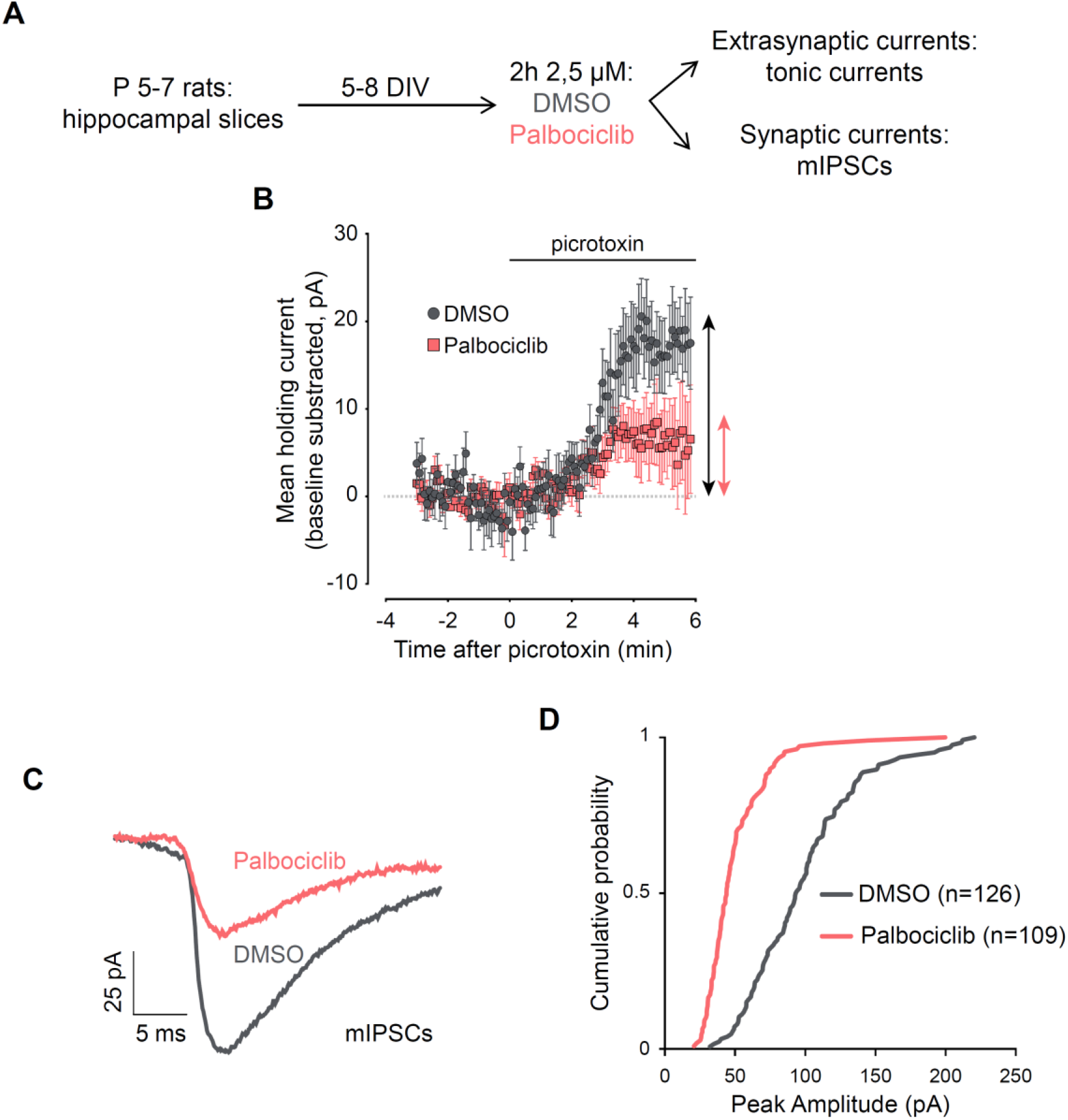
Cdk4/6 inhibition by Palbociclib decreases GABA tonic currents and mIPSC amplitude in rat hippocampal slices. A) Schematic representation of the experimental design for GABA currents recordings in CA1 neurons: hippocampal slices from newborn rats were cultured in vitro, treated for 2 h with DMSO or Palbociclib 2.5 μM, and tonic (extrasynaptic) or mIPSCs (synaptic currents) were measured. B) GABA inhibitory postsynaptic currents were evoked by stimulation of Schaffer collaterals and the tonic GABA_A_ receptor-mediated current was measured as the outward shift in holding current following application of the GABA_A_R antagonist picrotoxin. CA1 pyramidal neurons were recorded under whole-cell voltage-clamp at -60 mV. Effects of picrotoxin (100 μM) on holding currents recorded in DMSO-(grey rounds) or Palbociclib-treated (red squares) hippocampal slices are represented: the shift in holding currents is shown as double-headed arrows for DMSO (grey) and Palbociclib (red). Recordings are aligned at the time of picrotoxin addition (t=0). Each point represents the mean value of holding current at a specific time for 10 cells. C) To measure miniature inhibitory postsynaptic currents (mIPSC), 1 μM tetrodotoxin was added to block action potentials. Representative traces of mIPSC of DMSO- or Palbociclib-treated cells are shown. E) Cumulative distribution of mIPSC amplitude from CA1 pyramidal neurons treated with either Palbociclib or control vehicle, as indicated. “n” represents number of miniature responses recorded from nine (DMSO) or eight (Palbociclib) cells. (*p<0.001, Kolmogorov-Smirnov test).

**S3 Figure.**
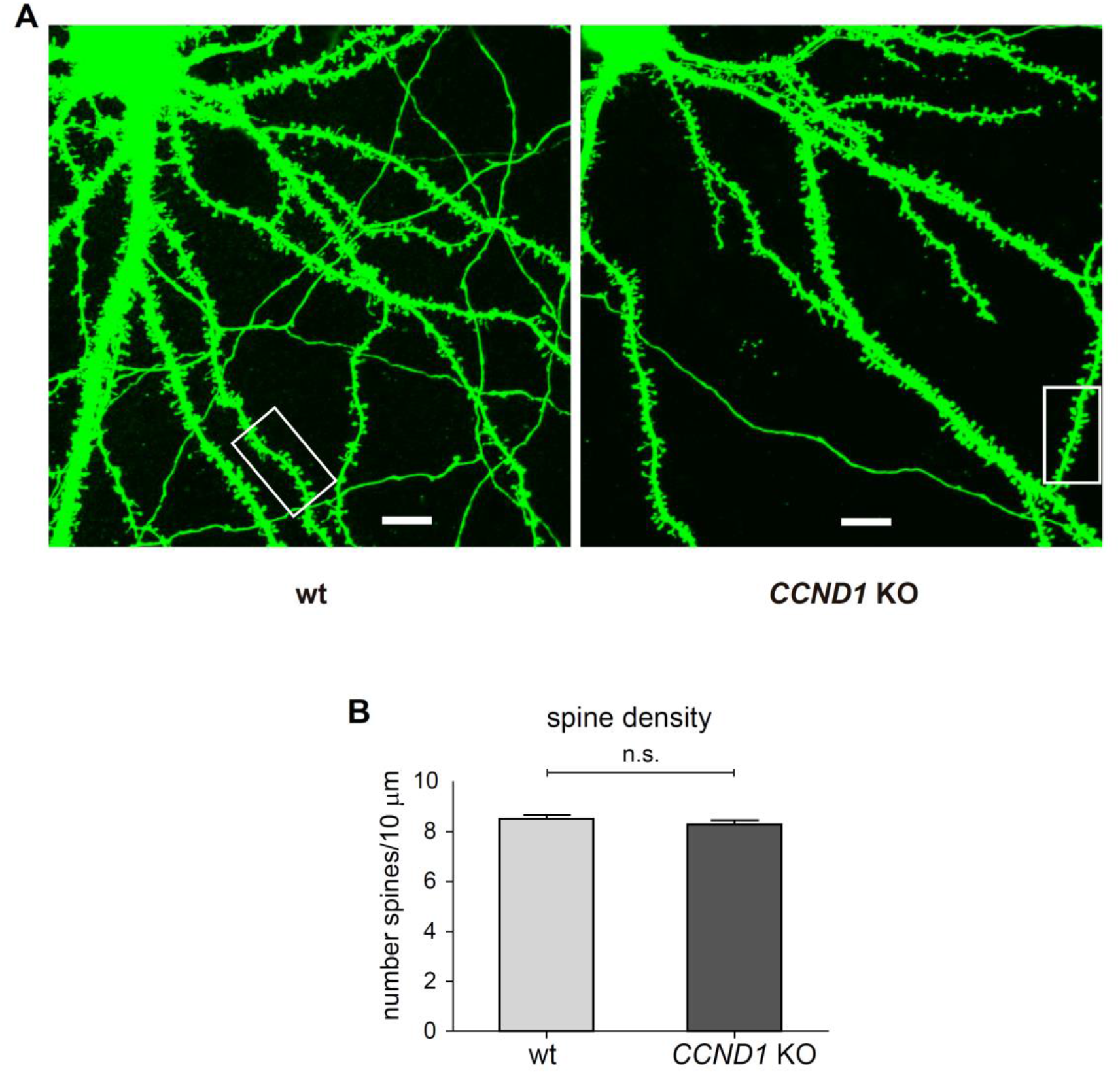
CCND1 KO increases the maturity of dendritic spines in cortical neurons. A) Dendritic spine analysis of WT and CCND1 KO mice cortical neurons at 20 DIV. Cultured cortical neurons were transfected at 3 DIV with GFP and fixed at 20 DIV. Representative confocal images of WT and CCND1 KO neurons. Scale bar: 10 μm. Changes in spine density and morphology were assessed by analyzing GFP-positive neurons with NeuronJ (see Materials and Methods). B) Quantification of the number of total dendritic spines relative to 10 μm of neurite, from WT (n= 59 neurons) and CCND1 KO (n= 62) neurons (4 mice/group). Bars show mean values ± S.E.M. ns: non-significant, Unpaired student’s t-test.

